# Variations of neuronal properties in the region of locus coeruleus of mice

**DOI:** 10.1101/2022.08.19.504582

**Authors:** Lucas Silva Tortorelli, Machhindra Garad, Marine Megemont, Sachiko Haga-Yamanaka, Anubhuti Goel, Hongdian Yang

## Abstract

Neurons in the locus coeruleus (LC) have been traditionally viewed as a homogenous population. Recent studies begin to reveal their heterogeneity at multiple levels, ranging from molecular compositions to projection targets. To further uncover variations of neuronal properties in the LC, we took a genetic-based tagging approach to identify these neurons. Our data revealed diverse spike waveforms among neurons in the LC region, including a considerable fraction of narrow-spiking units. While all wide-spiking units possessed the regular waveform polarity (negative-positive deflection), the narrow units can be further divided based on opposing waveform polarities. Under anesthesia, wide units emitted action potential at a higher rate than the narrow units. Under wakefulness, only one subtype of narrow units exhibited fast-spiking phenotype. These neurons also had long latencies to optogenetic stimulation. *In-situ* hybridization further supported the existence of a small population of putative GABAergic neurons in the LC core. Together, our data reveal characteristic differences among neurons in the LC region, and suggest that a fraction of electrophysiologically-identified narrow-spiking neurons can be fast-spiking interneurons, and their fast-spiking feature is masked by anesthesia.

## Introduction

The noradrenergic nucleus locus coeruleus (LC) has been implicated in a multitude of physiological and cognitive functions ranging from sleep-wake cycle and sensory perception to attention and memory (Berridge and Waterhouse, 2003; Aston-Jones and Cohen, 2005; Sara and Bouret, 2012; McBurney-Lin et al., 2019). Neurons in the LC have been traditionally viewed as a uniform population with homogenous intrinsic properties and functions (Aston-Jones and Bloom, 1981; Fallon and Loughlin, 1982; Loughlin et al., 1982; Waterhouse et al., 1993). One widely adopted criterion to identify LC neurons *in vivo* relies on their distinctive broad spike waveforms with prolonged after-hyperpolarization (e.g., (Andrade and Aghajanian, 1984; Aston-Jones et al., 1994; Berridge and Waterhouse, 2003; Rajkowski et al., 2004; Eschenko and Sara, 2008; Hirschberg et al., 2017)). Recent advances in single cell profiling and tracing (e.g., (Macosko et al., 2015; Shekhar et al., 2016; Tervo et al., 2016; Zingg et al., 2020)) have revealed heterogeneity in neuronal populations that were once thought to be homogeneous. These approaches have also begun to challenge the longstanding view of homogeneity in the LC (Chandler et al., 2019; Poe et al., 2020), uncovering heterogeneity at multiple levels including developmental origins, molecular phenotypes and projection targets (Robertson et al., 2013; Chandler et al., 2014; Schwarz et al., 2015; Kebschull et al., 2016; Kempadoo et al., 2016; Hirschberg et al., 2017; Plummer et al., 2017; Uematsu et al., 2017; Mulvey et al., 2018). Importantly, although narrow-spiking (sometimes neurochemically-defined GABAergic) neurons have been identified in the extended LC complex (Breton-Provencher and Sur, 2019; Kuo et al., 2020; Zouridis et al., 2024), it is believed that GABAergic interneurons do not exist in the LC core (Vreven et al., 2024). However, several electrophysiological studies that identified narrow-spiking neurons in the LC region were performed under anesthesia (Totah et al., 2018; Zouridis et al., 2024), and to our knowledge, it is currently unclear to what extent narrow-spiking, GABAergic interneurons exist in the LC core.

We took a genetics-based approach that allowed expressing ChannelRhodopsin-2 (ChR2) in noradrenergic (NA) neurons with a specific marker, dopamine-beta-hydroxylase (DBH), a key enzyme in the NA synthesis pathway downstream of dopamine. We recorded spiking activity from single units that were responsive to optogenetic stimulation. A subset of these units exhibited narrow spike waveforms and lacked the prolonged after-hyperpolarization. Interestingly, narrow-spiking units appeared to consist of two subgroups with opposing waveform polarities. These narrow-spiking units had lower spontaneous firing rate than the canonical wide-spiking units under anesthesia. Under wakefulness, only the group of narrow units with regular extracellular waveform polarity exhibited fast-spiking features. *In-situ* hybridization further revealed the existence of a small yet nonnegligible population of putative GABAergic interneurons in the LC core throughout the anterior-posterior axis. Together, this study presents new evidence to advance our understanding of the variations of neuronal properties in the LC region.

## Results

We implemented an approach to drive a genetically encoded ChannelRhodopsin-2 (ChR2(H134R)) transgene monoallelically from the *Gt(ROSA)26Sor* locus (*DBH-Cre/+;Ai32/+*). Taking this approach instead of using a viral vector should allow less variation in copy number and spatial spread when expressing ChR2 across DBH+ neurons. Because recent work showed that a subset of LC-NA neurons cannot be excited by short pulses (Hickey et al., 2014; Li et al., 2016), we used long pulses (0.2-0.3 s) to identify light-responsive neurons (Fig. 1a, b).

**Figure 1.**
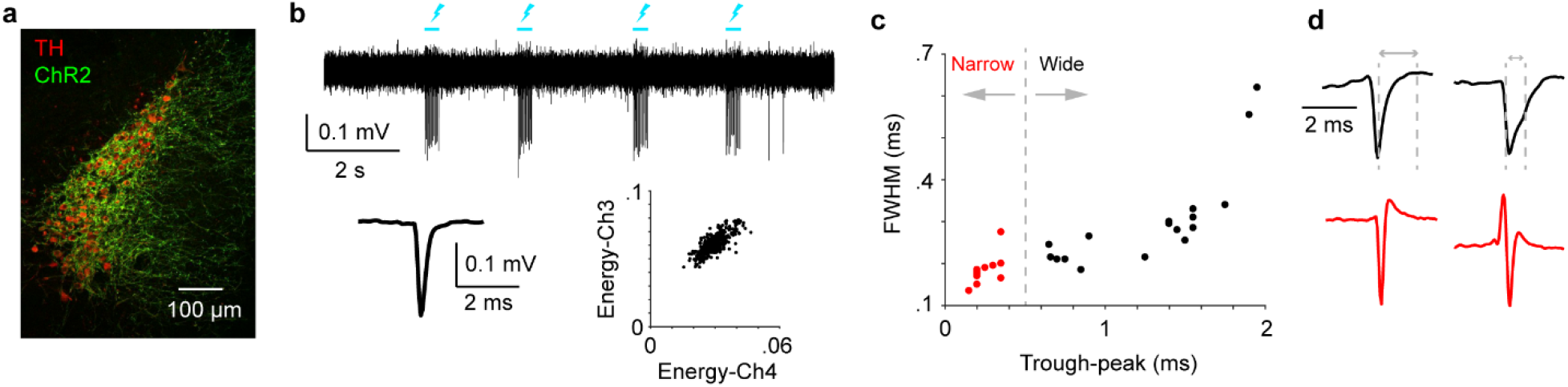
Putative LC units exhibit diverse waveforms. (a) ChR2 expression in a DBH-Cre/+;Ai32/+ mouse. TH: Tyrosine hydroxylase. (b)Top: Response of a ChR2-expressing LC neuron to optogenetic stimulation (lightning bolts). Bottom: Spike waveform and spike sorting diagram of this unit. (c)Scatter plot of trough-to-peak AHP vs. full width at half maximum (FWHM) for all units (n = 30). (d)Example waveforms for wide (black) and narrow (red) spiking units, and the illustration of trough to peak AHP and FWHM.

We first presented spike waveform quantifications among these neurons (30 units from 24 recordings of 19 mice). Typically, we were able to sort one single unit from a recording (20/24 recordings yielded 20 units), but a subset of recordings yielded multiple light-responsive units (4/24 recordings yielded 10 units). To ensure that all units were distinct with one another, one mouse only contributed one recording to the dataset, with one exception: when a second recording from the same animal was acquired on a separate day, and a light-responsive unit was sorted from a tetrode channel different from the first recording, the unit from the second recording was considered distinct from the first recording and included (a total of 5 units from 5 second recordings). We observed a spectrum of spike waveforms from this dataset (Fig. 1c). Similar to prior work (Totah et al., 2018), a threshold of 0.4-0.6 ms trough-to-peak after-hyperpolarization (AHP) robustly separated narrow from wide waveforms. A slight majority of these units were classified as wide (16/30), and the remaining as narrow (14/30). All wide-spiking units exhibited the typical extracellular waveform polarity (negative-positive deflection). However, narrow-spiking units also exhibited the reversed polarity (9/14, positive-negative deflection, Fig. 1d).

The presence of narrow units in/around the LC is consistent with recent work (Totah et al., 2018; Zouridis et al., 2024). Under anesthesia (2% isoflurane), all units emitted action potentials at a low rate (1.36 ± 0.36 spikes/s, Fig. 2a). However, the spontaneous firing rate of wide-spiking units was higher than the narrow-spiking units (Fig. 2b, wide vs. narrow, 2.12 ± 0.59 vs. 0.48 ± 0.22 spikes/s, P = 0.0029). This difference held when we further divided the narrow units into two subgroups based on waveform polarity (narrow-regular and narrow-reversed, Fig. 2c). These results suggest variable excitability among neurons in the LC region.

**Figure 2.**
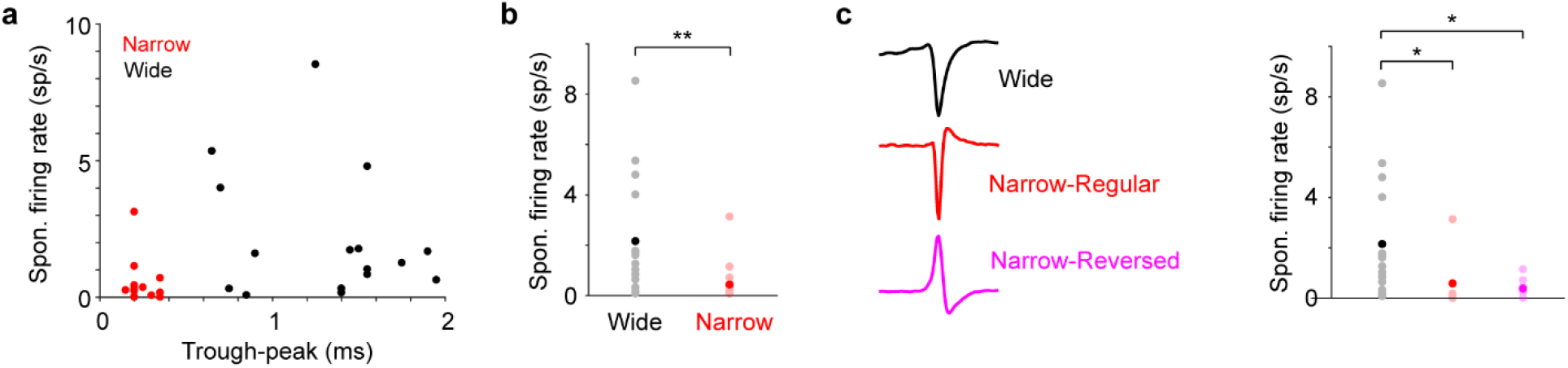
Wide and narrow units show different excitability under anesthesia. (a) Scatter plot of spontaneous firing rate vs. trough-to-peak AHP for all units. (b) Comparison of spontaneous firing rate between wide (n = 16) and narrow (n = 14) units under anesthesia with mean indicated. Wide vs. Narrow, 2.12 ± 0.59 vs. 0.48 ± 0.22 spikes/s, P = 0.0029. (c) Comparison of spontaneous firing rate between wide (2.12 ± 0.59 spikes/s, n = 16), regular narrow (0.67 ± 0.61 spikes/s, n = 5), and reversed narrow units (0.38 ± 0.12 spikes/s, n = 9) under anesthesia. Wide vs. Regular narrow, P = 0.028; Wide vs. Reversed narrow, P = 0.01; Regular narrow vs. Reversed narrow, P = 0.22.

Recent work identified fast-spiking interneurons in close proximity to the LC (Breton-Provencher and Sur, 2019). Could the narrow units we identified be fast-spiking? Since anesthesia can mask the fast-spiking feature (Patel et al., 1999; Zhao et al., 2021), this needs to be tested in the awake condition. In a subset of experiments, consecutive recordings were performed in the awake state following anesthesia (< 4 hours apart. 18 units recorded in the awake state: 8 wide, 5 regular narrow, 5 reversed narrow). Not surprisingly, the spontaneous firing rate of these units was higher in wakefulness than in anesthesia (Anesthetized vs. awake, 1.35 ± 0.36 vs. 10.73 ± 3.83 spikes/s, P = 2.6e-5), and this trend held for all three subgroups (wide, narrow-regular, narrow-reversed, Fig. 3a-c). However, only the regular narrow-spiking units exhibited >20Hz spontaneous firing rate in the awake state (27.89 ± 11.39 spikes/s, Fig. 3b). We noticed that for each of the 5 regular narrow-spiking units, spiking activity was consistently sorted from the exact same tetrode channel in both the anesthetized and awake states. Further analysis revealed high similarity between the waveforms sorted in the two states (waveform similarity 0.98 ± 0.007, Supp. Fig. 1), strongly suggesting that these units were tracked across time (Schoonover et al., 2021; Megemont et al., 2022), rather than that we randomly recorded a different set of fast-spiking neurons in the awake state.

**Figure 3.**
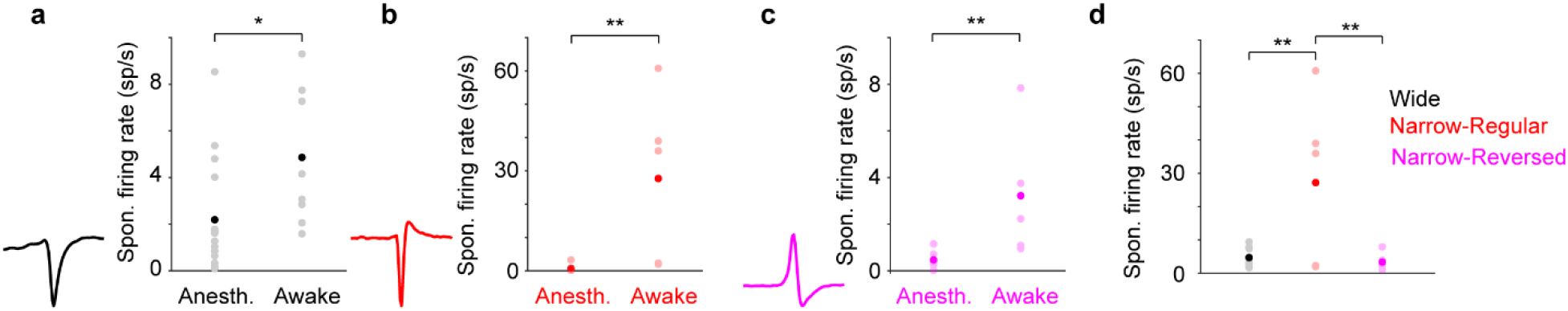
Neuronal excitability increases under wakefulness. (a) Comparison of spontaneous firing for wide units during anesthesia (n = 16) and wakefulness (n = 8). Anesthetized vs. awake, 2.12 ± 0.43 vs. 4.74 ± 1.04 spikes/s, P = 0.018. (b) Comparison of spontaneous firing for regular narrow units during anesthesia (n = 5) and wakefulness (n = 5). Anesthetized vs. awake, 0.67 ± 0.61 spikes vs. 27.89 ± 11.39 spikes/s, P = 0.001. (c) Comparison of spontaneous firing rate for reversed narrow units during anesthesia (n = 9) and wakefulness (n = 5). Anesthetized vs. awake, 0.38 ± 0.12 spikes vs. 3.16 ± 2.84 spikes/s, P = 0.001. (d) In the awake condition, the spontaneous firing rate of regular narrow units was higher than the wide units as well as the reversed polarity narrow units. Regular narrow vs. Wide, P = 0.001; Regular narrow vs. Reversed narrow, P = 0.005; Wide vs. Reversed narrow, P = 0.098.

The spontaneous firing rate of the regular narrow units was higher than both the wide units and the reversed narrow units (Fig. 3d). None of the reversed narrow units (0/9) showed fast-spiking phenotype, and as a group their activity was not different from the wide units in wakefulness (Fig. 3d). In addition, only the regular narrow units exhibited considerably longer latency to optogenetic stimulation than the wide units (Supp. Fig. 2), suggesting indirect, multi-synaptic excitation. Together, our data suggest that the narrow-spiking units with regular waveform polarity form a separate cluster, and are likely to be fast-spiking GABAergic neurons. It is known that GABAergic neurons are abundant just outside of the LC core (Jin et al., 2016; Breton-Provencher and Sur, 2019; Kuo et al., 2020; Luskin et al., 2023). While our histological confirmation did not have the spatial resolution to precisely pinpoint to what extent the recording sites were in the LC core, we did not find gross differences between where wide and narrow units were acquired (Supp. Fig. 3), suggesting that the narrow-spiking units in our dataset was not a result of systematic off-targeted recordings.

Since our electrophysiology data did not provide a definitive answer, we sought to further assess the existence of GABAergic interneurons in the LC core, and performed RNAscope *in-situ* hybridization (ISH) targeting the mRNA of glutamic decarboxylase 2 (GAD2), a marker for GABAergic neurons, alongside DBH (Methods). Our data revealed widely distributed GAD2 expression in areas close to the LC across the anterior-posterior axis. Importantly, neurons with significant GAD2 expression were found not only outside the LC core or along the LC boundary, but a small yet nonnegligible population was identified within the core (Fig. 4). To quantify GAD2-expressing neurons in the LC core, we first conservatively estimated the core boundary based on DBH expression. Next, given the varying levels of GAD2 expression, we established two conservative thresholds to define GAD2-expressing neurons (moderate expression: GAD2 signals within the soma is at least 3 sd above the background where virtually no GAD2 expression was found; high expression: GAD2 signals within the soma is at least 10 sd above the background. Fig. 4a, Supp. Fig. 4). Following this method, we identified a small population of high GAD2-expressing neurons throughout the anterior-posterior axis of the LC core (1.72 ± 0.32%, Fig. 4b, high GAD2). Including neurons with moderate GAD2 expression revealed a higher fraction within the LC core (5.51 ± 0.49%, Fig. 4b, all GAD2).

**Figure 4.**
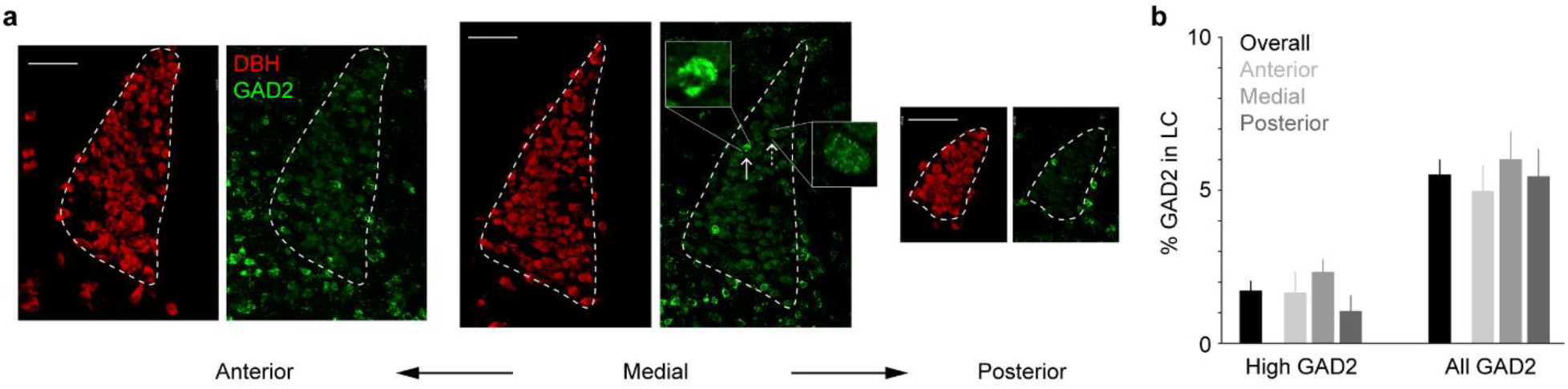
GABAergic neurons in the LC region. (a) Representative ISH (RNAscope) images illustrating DBH and GAD2 expressions in anterior, medial and posterior sections of the LC. Dashed lines illustrate the estimated LC boundary based on DBH expression. In the medial section, arrows and insets show a high GAD2-expressing neuron and moderate GAD2-expressing neuron. All images are of the same spatial scale. Scalebars: 100 µm. (b) Quantifications of the percentage of GAD2-expressing neurons in the LC core (5 anterior sections, 6 medial sections and 5 posterior sections from 3 mice). Percent of high GAD2-expressing neurons in the LC core: overall, 1.72 ± 0.32%; anterior, 1.66 ± 0.69%; medial, 2.33 ± 0.41%; posterior, 1.05 ± 0.52%. Percent of all GAD2-expressing neurons in the LC core: overall, 5.51 ± 0.49%; anterior, 4.98 ± 0.84%; medial, 6.01 ± 0.91%; posterior, 5.46 ± 0.88%.

## Discussion

Our data revealed diverse spike waveforms among neurons in the LC region, including a number of narrow-spiking units, consistent with recent work (Totah et al., 2018; Su and Cohen, 2022; Zouridis et al., 2024). While all wide-spiking units had the regular waveform polarity (negative-positive), the narrow-spiking units can be further divided into two subgroups: regular polarity and reversed polarity (positive-negative). Under anesthesia, wide-spiking units showed higher excitability as they had higher spontaneous firing rate than the narrow-spiking units. This trend was opposite in urethane-anesthetized rats, where narrow units had higher firing rate than the wide ones (Totah et al., 2018). However, a recent juxtacellular study reported that IHC-confirmed TH+ neurons exhibited higher firing rate than narrow-spiking TH-neurons under anesthesia (Zouridis et al., 2024). Current evidence suggests that adrenergic receptors are not primary targets of isoflurane (Schwinn et al., 1990; Schotten et al., 1998; Ranft et al., 2004; Ying et al., 2009), thus it is not likely that the different firing rates were due to isoflurane acting on subpopulations expressing different adrenergic receptors. Together with the evidence of their overall long response latency to optogenetic stimulation, at least a fraction of the narrow units identified in our dataset are likely to be TH-/DBH-.

All units increased spontaneous firing rate from anesthesia to wakefulness, and yet, only narrow-spiking units with regular waveform polarity exhibited fast-spiking phenotype, suggesting a subgroup of the narrow-spiking neurons in the LC region are fast-spiking GABAergic interneurons. The presence of GABAergic neurons in close proximity to the LC or along the rostral part of LC have been reported (Breton-Provencher and Sur, 2019; Luskin et al., 2023), even though the boundaries between the rostral part of LC and sub-coeruleus can be less well defined (Vreven et al., 2024). Admittedly, our electrophysiological results may be deemed circumstantial due to the limited spatial resolution to pinpoint the exact recording site. However, the recent juxtacellular study also identified a fraction of TH-, narrow-spiking neurons within the LC core (Zouridis et al., 2024). Furthermore, our *in-situ* hybridization data provide direct evidence of the presence of GAD2 mRNA at single-molecule level in the LC core, almost unequivocally demonstrating the existence of a small but nonnegligible group of GABAergic interneurons within the commonly considered LC boundary. A small number of inhibitory neurons can exert network-wide effects (Breton-Provencher and Sur, 2019; Kuo et al., 2020), yet investigating the functions of this sparse neuronal population poses a great technical challenge and awaits future work.

Our results may not seem surprising given that a minority of neurons in the LC core are known to be not noradrenergic and that LC neurons are known to produce multiple neurochemicals (e.g., (Berridge and Waterhouse, 2003; Mulvey et al., 2018)). However, our data suggest that without access to cell identity, caution should be exercised when categorizing narrow-spiking units in the LC core as noradrenergic and non-GABAergic, in particular under anesthesia when the fast-spiking phenotype can be masked (Patel et al., 1999; Zhao et al., 2021). These neurons could be interconnected with noradrenergic neurons, thus also respond to adrenergic pharmacology. Optogenetic tagging with short latency will greatly enhance the confidence of determining the identities of these neurons (as in (Su and Cohen, 2022)).

How about the narrow-spiking units with reversed waveform polarity? It is certainly possible that they consist of another subtype, given their distinct waveform and lower excitability compared to the wide-spiking units. On the other hand, experimental studies have suggested that narrow extracellular waveforms with a notable initial positive phase can originate from action potentials propagating along the axon (Raastad and Shepherd, 2003; Bakkum et al., 2013; Barry, 2015). Modeling work also supports that when the recording site is in the distal axonal compartment of a neuron, waveform can change and its polarity can flip (Gold et al., 2006). This would render spike waveform estimate inaccurate, categorizing the somewhat distorted axonal spike waveforms as narrow.

We note that not all wide-spiking units that were light-responsive had short latency to optogenetic stimulation. It has been reported that a considerable fraction (30-40%) of ChR2-expressing LC noradrenergic neurons can only be activated with prolonged illumination both *in vitro* and *in vivo* (> 100 ms, (Hickey et al., 2014; Li et al., 2016)). Altogether, these lines of evidence suggest heterogeneous excitability among LC noradrenergic neurons. However, we cannot exclude the possibility that long response latency on the order of tens of milliseconds is a result of indirect activation in a network with recurrent connections (Roux et al., 2014; Zouridis et al., 2024). Juxtacellular recordings combined with opto-tagging and cell filling (as in (Ding et al., 2022)) will be valuable to address this question.

One limitation of the current study is the relatively small sample size, which prevents further characterizing within-subtype properties (Chandler et al., 2014; Totah et al., 2018; McKinney et al., 2023). However, this is attributed to our efforts to avoid double counting, ensuring that each included unit is unique. As a result, this population should form a less biased representation in the LC region. Chronically implanted tetrodes have the potential to track putative identical units over time, allowing to assess their properties across different states.

## Materials and Methods

All procedures were performed in accordance with protocols approved by UC Riverside Animal Care and Use Committee (AUP 20190031). Mice were DBH-Cre (B6.FVB(Cg)-Tg(Dbh-cre) KH212Gsat/Mmucd, 036778-UCD, MMRRC); Ai32 (RCL-ChR2(H134R)/EYFP, 024109, JAX), singly housed in a vivarium with reverse light-dark cycle (9a-9p). 19 male and female mice of 8-12 weeks were implanted with titanium head posts as described previously (Yang et al., 2015). Custom microdrives with eight tetrodes and an optic fiber (0.39 NA, 200 um core) were built and implanted in the left LC to make extracellular recordings (Yang et al., 2021; Megemont et al., 2022). At the conclusion of the experiments, brains were perfused with PBS followed by 4% PFA, post-fixed overnight, then cut into 100 μm coronal sections and stained with anti-Tyrosine Hydroxylase (TH) antibody (Thermo-Fisher OPA1-04050).

Optogenetic stimulation for tagging consisted of a train of 200 or 300-ms pulses delivered at 0.3-0.5 Hz and 10 mW (RMS, measured at the tip of optical fiber). Tagging was performed during anesthesia (2% isoflurane) and sometimes during awake, non-task performing condition. Spike sorting was performed using MClust (Redish, 2014). 30 distinct single units (clustering quality measure, L_ratio_: 0.005 ± 0.002) from 24 recordings of 19 mice were included.

Data were reported as mean ± SEM unless otherwise noted. We did not use statistical methods to predetermine sample sizes. Sample sizes are similar to those reported in the field. We assigned mice to experimental groups arbitrarily, without randomization or blinding. We used two-tailed rank sum test except for small sample size such as regular narrow-spiking units (n = 5), in which case permutation test was performed.

The *in-situ* detection of DBH and GAD2 was carried out using the RNAScope (Advanced Cell Diagnostics), following our recent work (Nguyen et al., 2020). 3 mice were used for this analysis. Briefly, 20 μm brainstem slices were cut (Leica CM3050S Cryostat) and dried at 60°C for 30 min. Tissue sections were subsequently deparaffinized with 4% paraformaldehyde at 4°C (15 min), 50%, 70% and absolute ethanol (each 5 min) at room temperature (RT), followed by a 10-minute incubation with hydrogen peroxide. Epitope retrieval was accomplished by exposing the slides to RNAscope 1x Target Retrieval Reagent for 5 min at 99°C (98-102°C). Tissues were then permeabilized using RNAscope protease plus (ACD) for 30 min at 40°C, followed by a 2-hour incubation with specific Dbh and GAD2 RNA probes at 40°C. Dbh (ACD, 407851-C3), and GAD2 (ACD, 439371-C2) probes generated were uniquely amplified with Opal dyes (Akoya Biosciences, opal 570 (FP1488001KT), opal 650 (FP1496001KT)). Subsequently, Vectashield DAPI (Vector Laboratories H-1200) was used to counterstain tissue slides with DAPI and images taken on a Leica SPE confocal microscope. Fluorescent intensity was measured by Fiji software.

## Author contributions

LST and HY planned the project. LST and MG performed experiments. MG and HY analyzed data with assistance from AG and SHY. MG and HY wrote the manuscript with contributions from all authors.

## Acknowledgements

We thank Viji Santhakumar, Deepak Subramanian, Edward Zagha and Garret Anderson for comments on the manuscript. HY was supported by Klingenstein-Simons Fellowship Awards in Neuroscience, and NIH grants (R01NS107355, R01NS112200).

