## Supplementary figures for "Variations of neuronal properties in the region of locus coeruleus of mice"

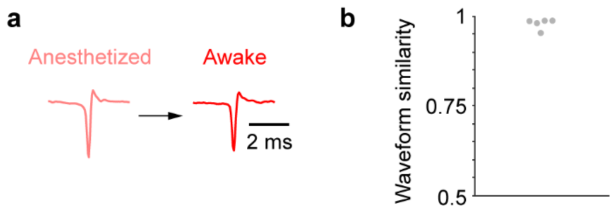


**Supp. Fig. 1**
(a) Example waveforms of a putatively identical narrow-spiking unit acquired in the anesthetized state and awake state.
(b) Waveform similarity measured as the Pearson correlation coefficient between spike waveforms during anesthesia and wakefulness for all narrow-spiking units with the regular polarity (0.98 ± 0.007, mean ± SEM, n = 5).

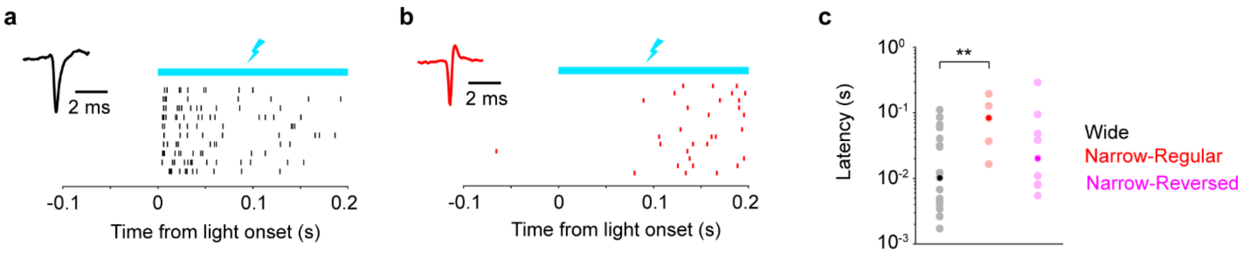


**Supp. Fig. 2**

(a) Example spike trains (ticks) to optogenetic stimulation of a wide unit, with 5 ms response latency to stimulation onset. Rows represent trials.

(b) Example spike trains (ticks) to optogenetic stimulation of a regular narrow-spiking unit, with 127 ms response latency to stimulation onset. Rows represent trials.

(c) Latency to optical stimulation in wide (n = 16), narrow-regular (n = 5), and narrow-reversed units (n = 9). Note that y axis is on logarithmic scale. Only the comparison between wide units and narrow-regular units revealed a difference, P = 0.0095.

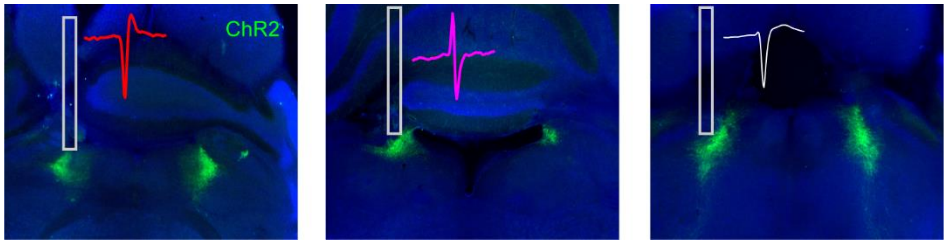


**Supp. Fig. 3**

Representative histological sections illustrating the estimated recording sites for acquiring narrow-regular polarity, narrow-reversed polarity and wide units.

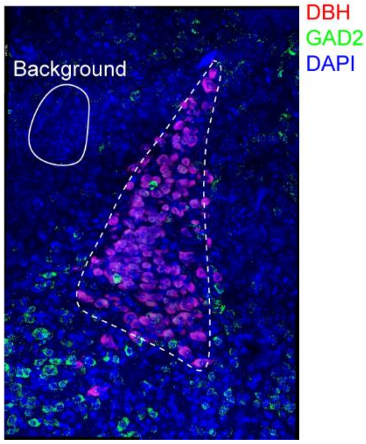


**Supp. Fig. 4**
Overlay of DBH, GAD2 and DAPI and the selected region with dense DAPI signals to quantify background GAD2 intensity.
